# M-Band Wavelet-Based Imputation of scRNA-seq Matrix and Multi-view Clustering of Cell

**DOI:** 10.1101/2022.12.05.519090

**Authors:** Zihuan Liu, Tong Liu, Wenke Sun, Yongzhong Zhao, Xiaodi Wang

## Abstract

Wavelet analysis has been recognized as a cutting-edge and promising tool in the fields of signal processing and data analysis. However, application of wavelet-based method in single-cell RNA sequencing (scRNA-seq) data is little known. Here, we present M-band wavelet-based imputation of scRNA-seq matrix and multi-view clustering of cells (WIMC). We applied integration of M-band wavelet analysis and uniform manifold approximation and projection (UMAP) to a panel of single cell sequencing datasets by breaking up the data matrix into a trend (low frequency or low resolution) component and (*M*-1) fluctuation (high frequency or high resolution) components. We leverage a non-parametric wavelet-based imputation algorithm of sparse data that integrates M-band wavelet transform for recovering dropout events of scRNA-seq datasets. Our method is armed with multi-view clustering of cell types, identity, and functional states, enabling missing cell types visualization and new cell types discovery. Distinct to standard scRNA-seq workflow, our wavelet-based approach is a new addition to resolve the notorious chaotic sparsity of scRNA-seq matrix and to uncover rare cell types with a fine-resolution.

**Author summary:** We develop M-band wavelet-based imputation of scRNA-seq matrix and multi-view clustering of cells. Our new approach integrates M-band wavelet analysis and UMAP to a panel of single cell sequencing datasets via breaking up the data matrix into a trend (low frequency or low resolution) component and (*M* – 1) fluctuation (high frequency or high resolution) components. Our method enables us to efficiently impute sparse scRNA-seq data matrix and to examine multi-view clustering of cell types, identity, and functional states, potentializing missing cell types recovery, fine rare cell types discovery, as well as functional cell states exploration.

## Introduction

Recent breakthroughs in methodology are enabling the study of the transcriptomes of individual cells, which paves the way for more objective investigations of cellular functions at single-cell level [1–3]. However, the sparsity of droplet-based single-cell RNA sequencing (scRNA-seq) data, i.e., a large number of expressed genes with zero, unmeasured, unknown or low read counts (known as “dropout” events) being presented, as a formidable problem to be resolved, has been greatly compromising the performance of exploratory analysis of cell identities, types, and states, as well as the identification of respected gene signatures [4–6]. Thus, how to impute the scRNA-seq data matrix is crucial for scRNA-seq analysis. Imputations based on parametric modeling techniques incorporating with bimodal mixture distribution and multimodal distributions, such as SCDE [4], MAST [7], BPSC [8], DEsingle [9], Monocle [10], D3E [11] and scDD [12], have been largely constrained by the multiscale complex distributions of the measured gene expression of droplet-based scRNA-data. In addition, these methods are bounded by simplified predetermined parametric distribution assumptions, requiring the specification of a number of parameters and are incapable of dealing with more complicated situations.

An additional primary objective of scRNA-seq analysis is to identify and discover cell types [13–16], identities [17], states [15,18], as well as accompanied gene signatures [19, 20], while canonical pipelines have been unsatisfying due to limited resolution even with missed critical cell types, identity, and functional states. Earlier studies based on microscopy [21], histology [22], and pathological criteria [23] have contributed to the resolution of this problem. Recent efforts have utilized both unsupervised and supervised clustering techniques to cluster cells alongside transcription signatures [14, 16] yet with big room to be optimized, requesting novel approaches.

Those limitations in the existing methods have motivated us to develop a new method, aiming to resolve sparsity due to dropout events, recover missing cell clusters, and uncover rare cell types. Wavelet frameworks have been successfully applied to numerous tasks in the biomedical domain, such as genomics [24], promoters [25,26] and GWAS [27]. However, wavelet-based method in scRNA-seq data analysis has been largely unexplored.

Our hybrid technique incorporated with M-band orthogonal wavelet enables us to illustrate multi-view clustering of cell types, identities, and functional states from a variety of perspectives simultaneously. Distinct from the canonical RNA-Seq analysis pipelines, our strategy can help resolve the notorious chaotic sparsity of droplet RNA-Seq matrix and have the power to uncover missed and rare cell types, identities, states, via wavelet-based imputation with bandwidth selection and multi-view of clustering (WIMC). In particular, we propose a non-parametric wavelet-based imputation algorithm of sparse data that integrates M-band orthogonal wavelet for recovering dropout events of scRNA-seq datasets. For such imputation, we split the data matrix into *M* components, including a trend component (with low frequency or low resolution) and a (*M* – 1) fluctuation component (with high frequency or high resolution). The trend component of the M-band wavelet enables us to transform data automatically impute the original sparsity of the RNA-Seq matrix by taking a weighted-average like process.

## Results

### The overview of WIMC

We aim to build a pipeline with sparse scRNA-seq data imputation and multi-view of clustering. Stemming from the canonical scRNA-seq workflow, our WIMC integrates wavelet-based non-parametric imputation with filter length selection and multi-view of clustering into canonical pipelines. In the light of wavelet analysis, we develop WIMC to impute sparse data matrix followed by multi-views of scRNA-seq in different frequency windows, potentializing recovery of missing cell types, cell states, cell identities, as well as rare cell types discovery. Our WIMC consists of seven essential steps, including quality control, log-transformation, DWT, principal component analysis (PCA) dimension reduction, multi-view with uniform manifold approximation and projection (UMAP) based clusters visualization, gene signature and identification of cell types, identities, states, and performance assessment.

Given a raw scRNA-seq sparse data matrix (Fig.1A), we remove the low-quality cells (Fig.1B) to ensure that technical effect does not distort downstream analysis results. We next perform logarithm transformation of the raw sparse matrix (Fig.1C) followed by DWT with different filter banks and filter lengths on the logarithm transformed matrix (Fig.1D). This step allows us to impute spares data matrix (Fig.1E). The above efforts enable canonical data matrix decomposition into different frequency components, resulting in a multi-view clustering of cell types, identities, and functional states from a variety of perspectives simultaneously. In addition to developing our unique DWT, we apply PCA on both wavelet-based imputed and canonical matrices for dimension reduction. We next apply UMAP method for exploratory visualization (Fig.1G). We annotate clusters by default clustering method implemented in Seurat [28]. In particular, on PCA-reduced matrix, we implemented Louvain community detection algorithms [30] on single-cell K-Nearest Neighbor (KNN) graph to organize cells into clusters. To characterize the similarities and differences between classic and our proposed method, we compare the clusters based on the canonical and wavelet-based imputed multi-view matrices (Fig.1H). Of note, we used the single-cell Cluster-based automatic Annotation Toolkit for Cellular Heterogeneity (scCATCH) [29] for cluster annotation. Finally, to assess the performance of WIMC multi-view, we analyze the intersection of the marker genes and cell types resulting from canonical method and our WIMC approach.

**Fig 1.**
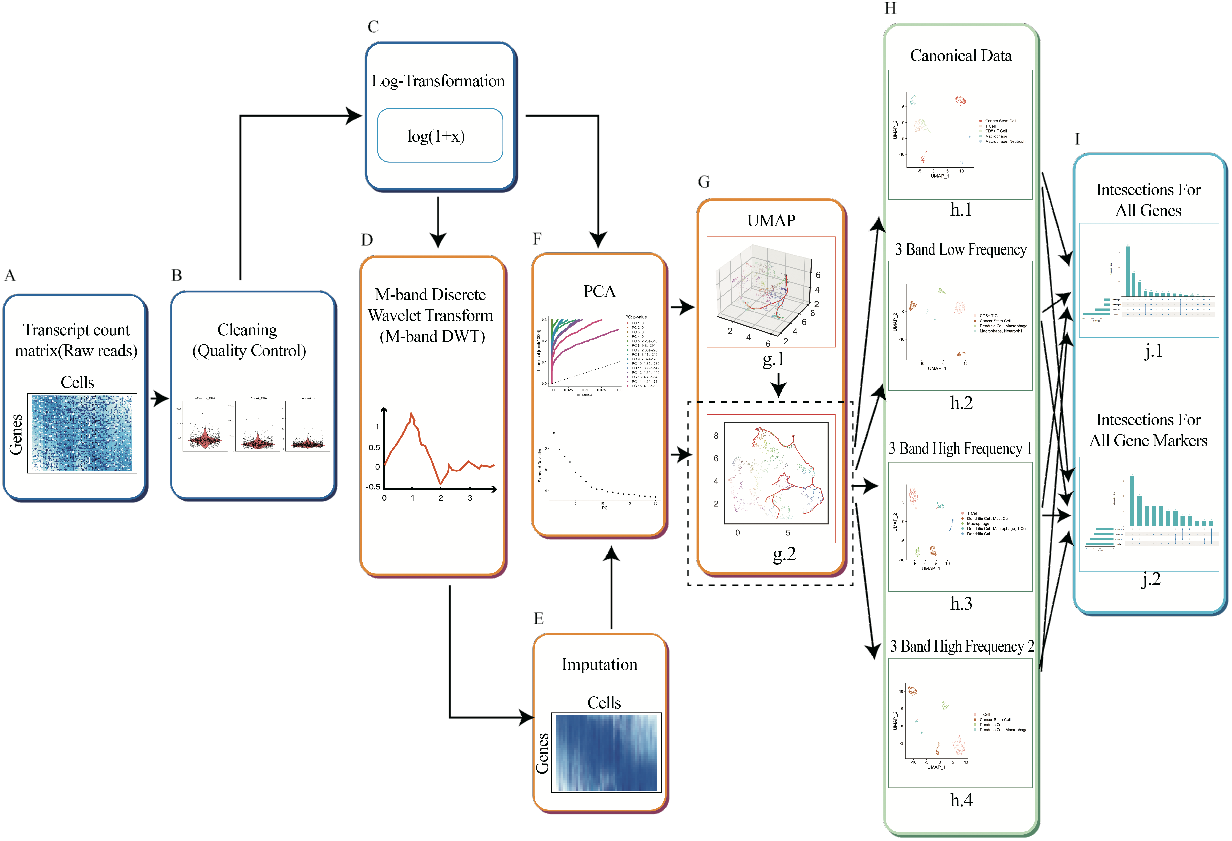
Illustration of the WIMC workflow. (**A**) Raw scRNA-seq sparse data matrix with row as transcripts and columns as individual cells. (**B**) Quality control diagrams, demonstrating the process of removing unqualified cells and transcripts involving mitochondrial genes. (**C**) Logarithm transformation of the raw sparse matrix. (**D**) Applying M-band DWT on the logarithm transformed matrix. (**E**) Imputation implemented via M-band DWT. (**F**) PCA-based orthogonal transformation and dimension reduction on wavelet-based imputed and canonical matrices. (**G**) UMAP visualization of both PCA-based canonical and wavelet-based imputed matrices. (**H**) Assessing the performance of WIMC.

### Mathematical statement of WIMC

Let 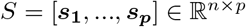 denotes the canonical data matrix and 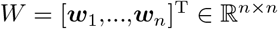 be the corresponding M-Band DWT matrix [31,32]. Then, the M-Band DWT of ***s**_i_* for *i* = 1, …,*p* is given by

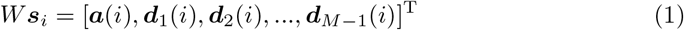

where *n* = *MK*, *a*(*i*) = [*a*_*i*1_, *a*_*i*2_, …, *a*_*ik*]^T^ and ***d**_j_*(*i*) = [*d*_*j, i*, 1_, *d*_j, i_, 2_, …, *d*_*j, i, k*_]^T^ for *j* = 1, 2,…, *M* – 1. For brevity, we denote *W**s**_i_* by 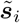. Let 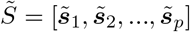, then

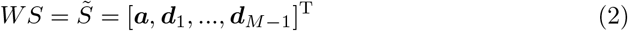

where ***a*** = [***a***(1), …***a***(*p*)] being the lowest frequency component (or trend) and ***d***_1_ = [***d***_1_(1),…, ***d***_1_(*p*)],…, ***d***_*M*−1_ = [***d***_*M*−1_(1),…, ***d***_*M*−1_(*p*)] being the higher frequency components (or fluctuations) of *S* in wavelet domain. Since *W* is an orthogonal matrix, {***w***_1_, ***w***_2_,…,***w**_n_*} forms an orthonormal basis of 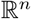. The components of 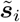 are also called the wavelet coefficients of ***s**_i_*. Therefore, the components of 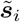 are coordinates of ***s**_i_* under this wavelet basis. Due to the orthonormality of *W*, for *i* = 1, 2, …, *p*, we have 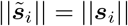 and *W*^T^ = *W*^-1^. Hence the M-Band DWT preserve the length or energy of vectors it transforms. If we multiply both sides of (2) by *W*^T^, we obtain

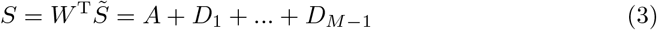

and

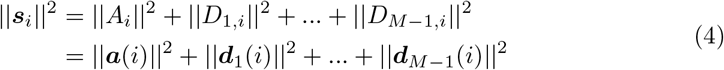

where *A* = [*A*_1_, *A*_2_,…, *A_P_*] = [***w***_1_, ***w***_2_,…, ***W**_k_*]***a*** and *D_j_* = [*D*_*j*1_, *D*_*j*2_, …, *D_jp_*] = [***w***_*kj*+1_, ***w***_*kj*+2_,…, ***w***_*kj*+*k*_]*d_j_*, *j* = 1, 2, …, *M* – 1. In other words, we use the M-Band DWT to decompose S into the sum of M orthogonal components (multi-view windows) with one low frequency component and *M* – 1 higher frequency components, as shown in (3)

We next aim to find *A_i_* and *D_j_* for *i* = 1, … *p* and *j* = 1, 2,…, *M* – 1. Define 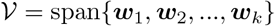, *i* = 1, 2,…, *M* – 1, then 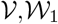, … and 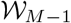 are orthogonal subspaces of 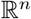. It follows that,

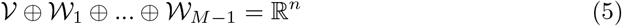

Let 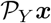 be a projection of ***x*** on *Y*, we have

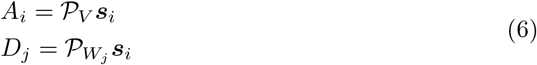

We then apply the UMAP on S and its corresponding components *A, D*_1_,…, *D*_*M*−1_ to obtain their 2*D* multi-view images as 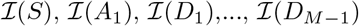, resulting in multi-view clusters with principal cell names, encompassing both low frequency and high frequency components of *S*. Thus, the essemble of annotated gene clusters enables us a multi-views cell types, identities, and states with sc-RNA seq data.

We assess the performance of WIMC via computing the intersection of 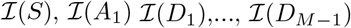. Since the space of different frequencies are orthogonal to each other, few genes could be found in the intersection of different components theoretically, Therefore, the smaller number of genes appearing in the intersection of these components yields a better resolution of clustering.

### Imputation of sc-RNA-seq sparse matrix

We first carry out WIMC imputation on a range of published scRNA-seq datasets, including breast cancer data (CID-3921, CID-4495, CID-4523 and CID-4463) [33], colorectal cancer data [34] and human peripheral blood mononuclear cells (PBMCs) from 10 × Genomics [35]. We summary the performance of WIMC (Table 1). On the one hand, it appears that the longer filter results in an increased proportion of non-zero elements than shorter filter. Oversized filters, on the other hand, might cause frequency components overlapping, which will be discussed in a later chapter. Specifically, for breast cancer data, the proportion of non-zero elements is approximately 28.29% by employing the Daub4 DWT, while canonical data matrix respective none sparsity of 5.94%; this can be further improved to 74.50% with longer filters. We find similar performance across other datasets.

**Table 1.**
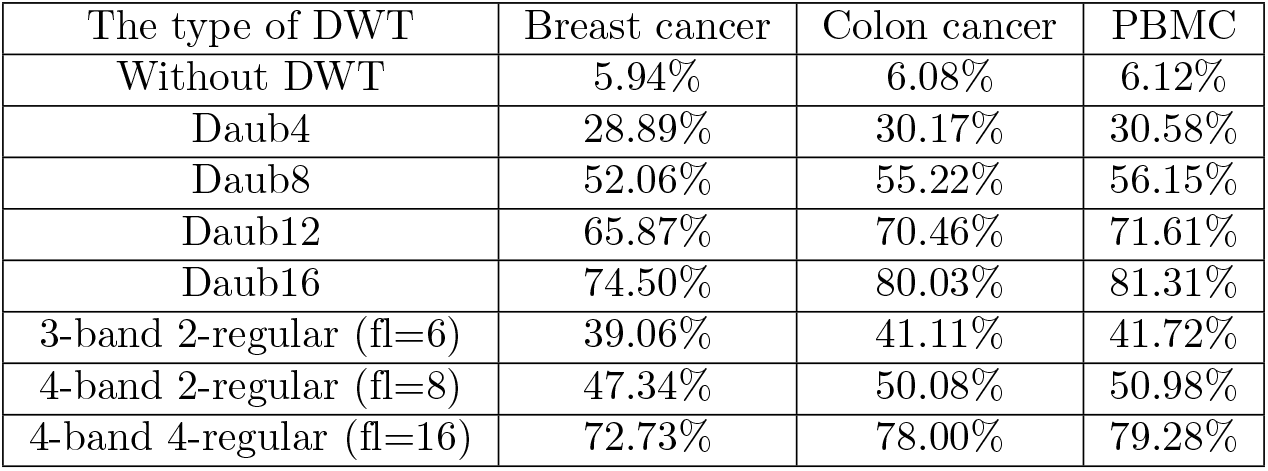
The percentage of non-zero elements in the count matrix before and after wavelet transform. Breast cancer refers to data CID-3921. Daub-2n, belonging to Daubechies wavelet transform family, refers to 2-band n-regular DWT with filter length(fl) being 2n.

### Cluster analysis of the scRNA-Seq dataset

Next, we visualize the representation of cell types (Fig.2) learned by the WIMC in the two-dimensional UMAP space with breast cancer data (CID-3921). It appears novel clusters to be emerging from 2-band (Fig.2B), 3-band (Fig.2C), and 4-band (Fig.2D) DWT in comparison with canonical UMAP clustering (Fig.2A). We use the 2-regular edition of M-band DWTs throughout this work unless otherwise specified. There are 8 cell types in the canonical UMAP setting, while there are 11, 17 and 16 types in 2-band, 3-band and 4-band UMAP settings, successively. In addition to distinguishing the cell clusters observed in the canonical method, including cancer stem cells, CD4, T cells, B cells, Hepler T cell, Dendritic cell, and Cytotoxic Cell, we find clusters of killer cells, eosinophil cells, and plasma cells with 2-band, 3-band and 4-band DWT, and smooth muscle cells with 3-band and 4-band DWT, as well as some unknown cell clusters with 3-band and 4-band DWT. It appears that novel clusters revealed by WIMC are missing cell types or rare cell types, yet to be validated. Consistently, with more breast cancer scRNA-seq data, WIMC helps discover missing cell types and rare cell types, further demonstrating its power in uncovering novel clusters (Fig.S1-S3).

**Fig 2.**
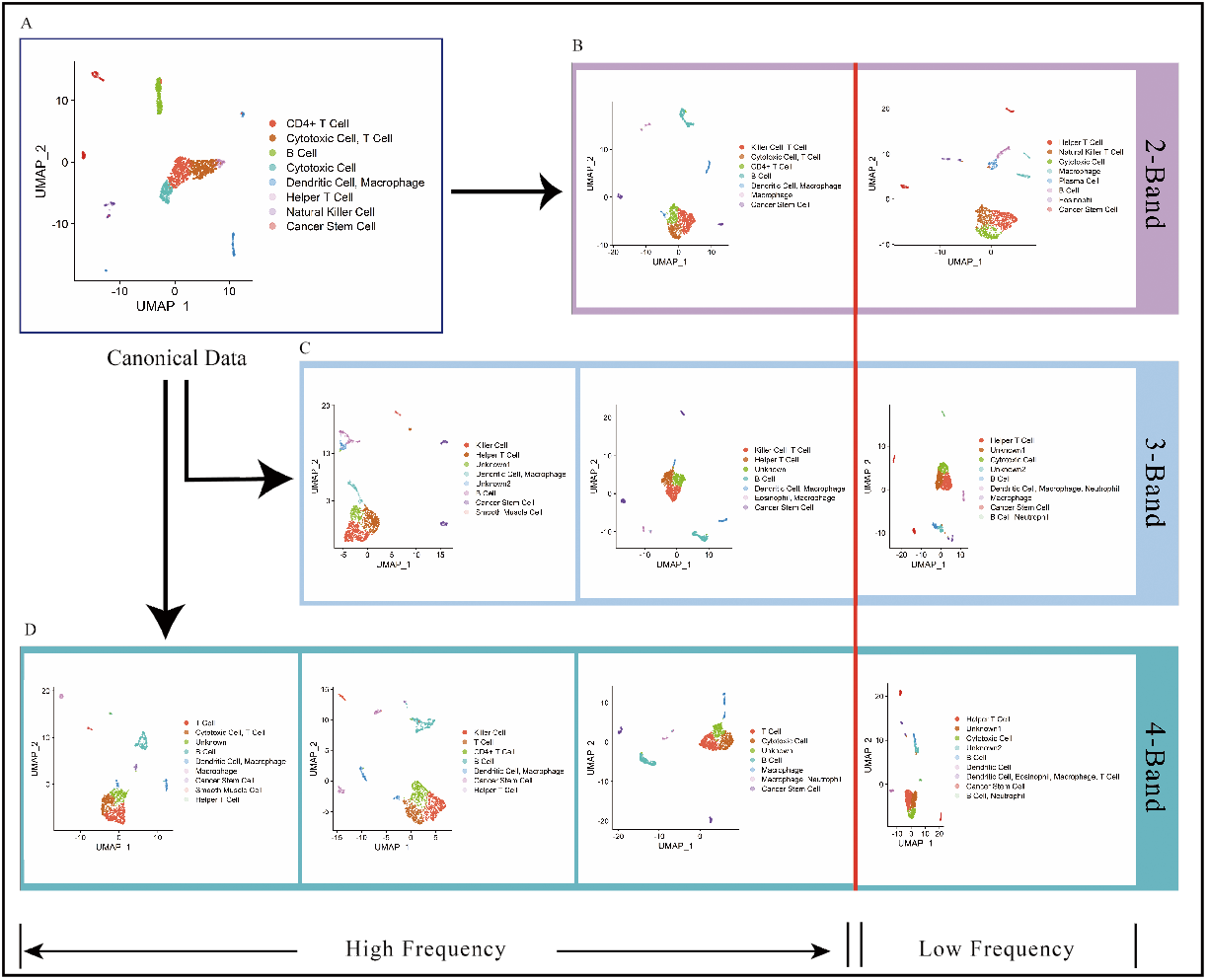
Multi-view of clusters of the breast cancer dataset. (**A**) UMAP visualization of cell types based on the canonical matrix with 8 types of cells (The number of cells in each cluster are 271, 240, 163, 151, 94, 56, 55 and 93). (**B**) Clusters based on the 2-band DWT, in which the low frequency component-based clusters contain 8 cell types (The number of cells in each cluster are 296, 229, 194, 91, 84, 83, 53 and 93) and high frequency component-based clusters contain 7 cell types (The number of cells in each cluster are 316, 212, 190, 168, 90, 54 and 93). (**C**) Clusters based on the 3-band DWT, where the low frequency component-based clusters contain 9 cell types (The number of cells in each cluster are 288, 241, 191, 95, 71, 60, 53, 93 and 31), high frequency 1 component-based cluster contain 7 cell types (The number of cells in each cluster are 283, 223, 207, 169, 94, 54 and 93) and high frequency 2 component-based cluster contain 8 cell types (The number of cells in each cluster are 303, 284, 149, 90, 65, 105, 93 and 34). (**D**) Clusters based on the 4-band DWT, in which the low frequency component-based clusters contain 9 cell types (The number of cells in each cluster are 289, 249, 181, 100, 67, 60, 53, 93 and 31), high frequency 1 component-based cluster contain 7 cell types (The number of cells in each cluster are 289, 243, 184, 169, 91, 54 and 93), high frequency 2 component-based cluster contain 7 cell types (The number of cells in each cluster are 315, 220, 214, 171, 91, 93 and 19) and high frequency 3 component-based cluster contain 9 cell types (The number of cells in each cluster are 308, 229, 179, 169, 90, 51, 42, 34 and 21).

We further access WIMC performance on a benchmark scRNA-seq experiment that profiled 2613 peripheral blood mononuclear cells (PBMCs) from 10 × Genomics. Specifically, we observe that WIMC discovers 8, 10 and 13 types of cells in 2-band, 3-band and 4-band UMAP settings, whereas canonical method reveals 6 cell types (Fig.S4). In addition to the six cell types which have been identified by the canonical method, WIMC discover new clusters of natural killer cells with 2-band and 3-band DWT, non-switched memory B cells with 3-band and 4-band DWT, regulatory T cells, memory T cells and SLC16A7+ cells with 4-band DWT, as well as some unknown cell types with 3-band and 4-band DWT. Again, those unknown cell types most likely are rare cell types that can be functionally validated in a biological experiment.

Moreover, we analyze our proposed method on a published scRNA-seq dataset from a colorectal cancer patient. A similar analysis can be performed on this dataset, as Fig. S2 illustrates that there are 6 types in the canonical UMAP setting and while there are 9, 12 and 12 cell types in 2-band, 3-band and 4-band UMAP settings, respectively. In addition to the cell clusters observed in the both canonical and WIMC method, our method identifies additional cell clusters, including Mitotic Fetal Germ cell, Astrocyte cell Monocyte cell, Fibroblast cell, Plasma cell and unknown cell types.

### Distinct gene signatures of cell clusters from different Wavelet-Based multi-view windows

To quantify the improvement in multi-view clustering of cells from our algorithms, we assess the performance of clustering at two levels, overlapping’s of cells between clusters and intersections of gene markers.

First, we compute pairwise p-value (Fig.3) with multi-test correction between different frequency components to determine if each intersection can be recognized as an independent cluster. With significance level of 0.01, we find 10 significant clusters using 2-band DWT for the breast cancer data (CID-3921), and one more cluster from high frequency component can be found when compared with canonical data. However, the effects are marginal when utilizing multi-band DWT. Of note, increasing the number of intersections between clusters in different frequency components may result in some subsets containing a small number of cells with a relatively high p-value. We discover 22 significant clusters using 3-band DWT, where 10, 8 and 11 clusters are found in low, high 1 and high 2 frequency component-based data, respectively. Similarly, a total of 28 significant clusters are revealed for 4-band DWT, with 10, 8, 9 and 9 clusters in low, high 1, high 2 and high 3 frequency component-based data, respectively (Fig.3 and see Fig.S6-S10 for more detailed results). Consequently, one can use WIMC to divide some clusters in canonical data into a few subgroups, resulting in the clusters with greater significance.

**Fig 3.**
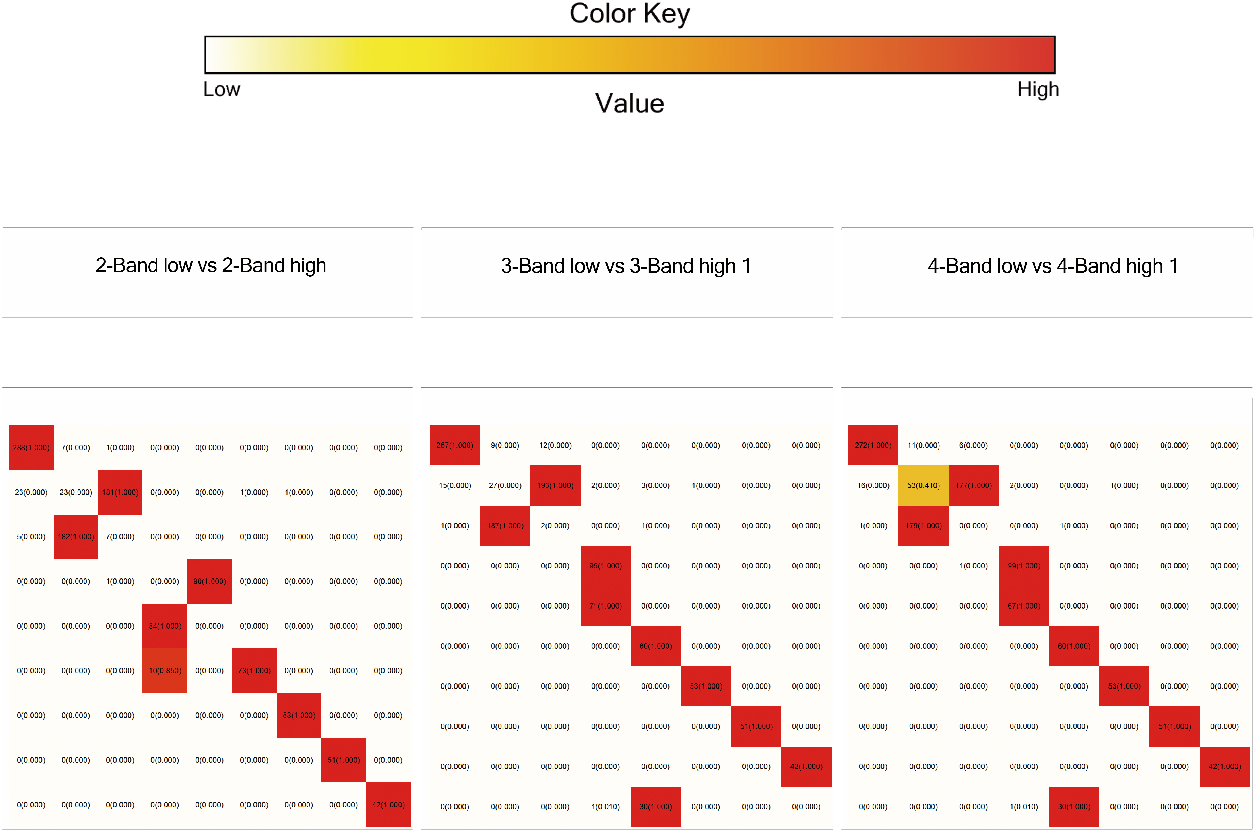
Heatmap of pair-wise intersection of clusters among different frequencies of 2-band, 3-band and 4-band DWT for the breast cancer dataset. Each cell in heatmap represents the number of cells (and the p-values) of corresponding intersection. The color key indicates the magnitude of the p-value. Dark hues indicating clusters that are more likely to be statistically significant, for light hues vice versa.

Apparently, the cell types in different frequency components can overlap or be distinct. Indeed, comprehensive intersection analysis of gene signatures further support the power of multi-view of WIMC (Fig.4).

**Fig 4.**
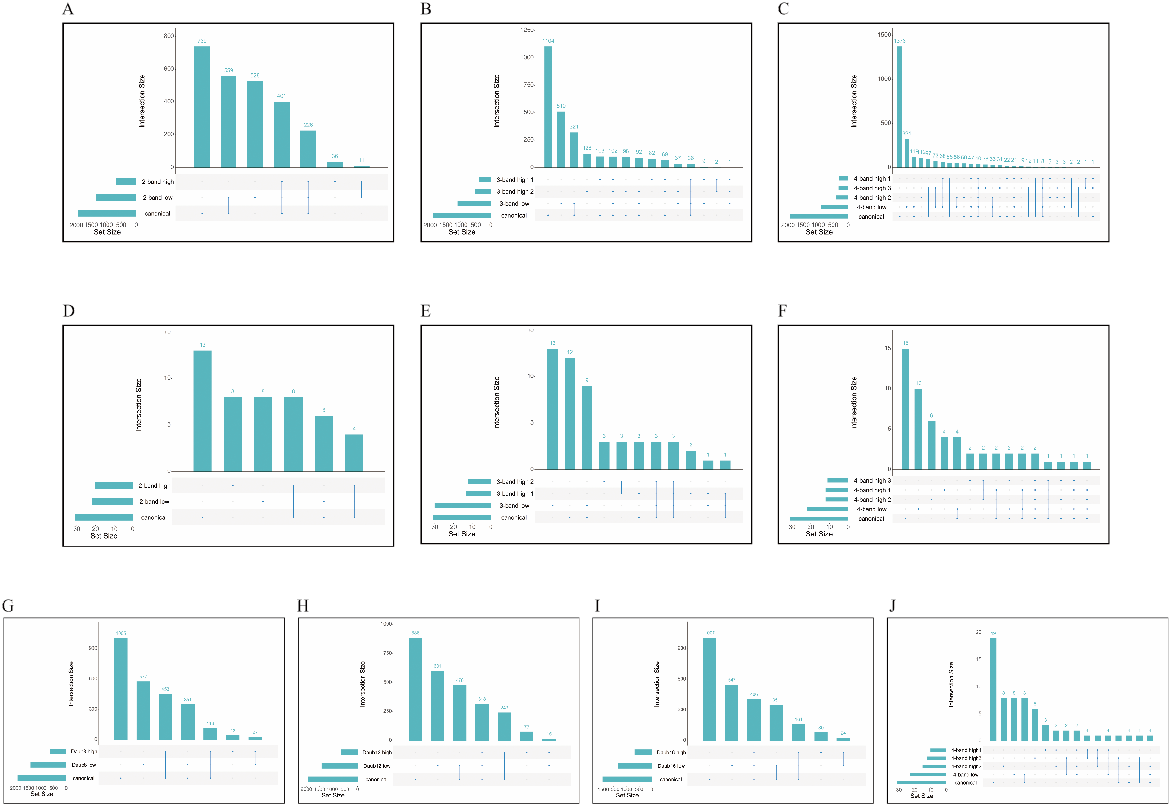
Assessing the performance of WIMC on the breast cancer dataset. The x-axis of panels (A) through (J) represents the total number of genes in each component and the y-axis of panels (A) through (J) shows the number of genes in each intersection set. The cyan dots represent the number of genes belonging to the corresponding components, for gray dots vice versa. (**A**) The intersection of genes between canonical data and its different frequency components using 2-band DWT (Daub4). (**B**) The intersection of genes between canonical data and its different frequency components using 3-band DWT. (**C**) The intersection of genes between canonical data and its different frequency components using 4-band DWT. (**D**) The intersection of cell-type related markers between canonical data and its different frequency components using 2-band DWT (Daub4). (**E**) The intersection of cell-type related markers between canonical data and its different frequency components using 3-band DWT. (**F**) The intersection of cell-type related markers between canonical data and its different frequency components using 4-band DWT. (**G**)) The intersection of genes between canonical data and its different frequency components using 2-band DWT (Daub8). (**H**) The intersection of genes between canonical data and its different frequency components using 2-band DWT (Daub12). (**I**) The intersection of genes between canonical data and its different frequency components using 2-band DWT (Daub16). (**J**) The intersection of cell-type related markers between canonical data and its different frequency components using 4-band 4-regular DWT.

Daub-2n DWT, which is 2-band n-regular DWT with filter length 2n, produced one low-frequency component and one high-frequency component. Low-frequency parts are traditionally consider as an approximation of canonical data and share a number of common genes with canonical data. However, we find 528 more genes in the 2-band low-frequency component-based data than in canonical data (Fig.4A). Similarity, we discover 510 additional genes in 3-band low-frequency component-based data compared to canonical data (Fig.4B), and 321 more genes in 4-band low-frequency component (Fig.4C) than in canonical data.

DWT has the ability to decompose canonical data into orthogonal frequency components, which is another important property. This means that different frequency parts are not overlapped theoretically. Using Daub4 WT, there are 237 genes that overlap in both frequency components, but only 11 (4.6%) of them are new, and the remaining 226 genes appeared in the canonical data (Fig.4A).

The high-frequency components play an important role in recognizing cell types. Compared with the canonical data, we identify more cell-type related gene markers in the high-frequency components under multi-band DWT (Fig.4D–4F). Different from the low frequency component, high frequency components describe the detail characteristics which may not be discovered under traditional process.

Similar phenomena can be found in colorectal and PBMC dataset. In PBMC dataset, there are 133 genes that are identified at the intersection of two frequency components using 2-band DWT, whereas only 2 genes are absent from canonical data. WIMC performs better on this data in finding cell-type related markers. Under 2-band, 3-band or 4-band DWT, there are 14, 16 and 20 new non-overlapped markers, respectively (Fig.S14). For the colorectal dataset, we only discover 80 at the intersection between two frequency components using 2-band DWT, which is significantly less than the number of new genes only appearing in one single component. For example, we discover 301 and 201 genes in 2-band low frequency component-based and high frequency component-based data (Fig.S15).

## Discussion

In this study, we develop WIMC, a M-band DWT-based approach for successful imputation of sparse scRNA-seq matrices and multi-view clustering of cells. WIMC not only offers a unique capability that overcomes key limitations of existing single-cell imputation techniques, but also helps to identify missing cell types and rare cell types alongside activity. First, by utilizing a non-parametric wavelet-based model, the proposed method does not require any predetermined parametric distribution assumptions, allowing our models to function in more complex situations. Second, our imputation techniques are broadly applicable to other types of missing data where imputation has been useful, such as time series data [**?**]. Third, WIMC can help discover rare cell types missed by canonical method with a fine resolution. Finally, WIMC can potentially prioritize rare cell types for experimental validation by incorporating intersection analysis on multi-view of clusters.

Our study is subject to several limitations. The length of filters has varying effects on imputation performance, leading to selection of length of filters challenging. Indeed, when compared with canonical data, we find 577, 601, and 547 more genes in low frequency components and 43, 77, and 80 more genes in high frequency components using Daub8, Daub12, and Daub16 DWT successively. (Fig.4G–4I). It is most likely that more marker genes can be found in high frequency components as the filter length increases, yet with some exceptions (Fig.S12-S13).

In addition, WIMC performs better on finding cell-type related markers with 4-band 4-regular DWT using filter length of 16 (Fig.4J). The high-frequency components still provide new gene markers, and we find less intersection between different frequency components. Daubechies DWT produces less overlapping between different frequencies, while M-band DWT with *M* ≥ 3 has the potential to discover more gene markers. However, given that optimized clusters must be physiology-relevant with biological significance, an optimized band number (M) might be required.

Lastly, even though multi-view clusters can detect unidentified cell types that may represent novel cell types in the biological field, these cell types are yet to functionally validate in a biological experiment. For future research, we envision that DWT-based framework integrated with other omics information is promising. Meanwhile, WIMC has a great potential to analysis spatial single-cell gene expression data. Thus, our WIMC paves the way to deploy wavelet tools in scRNA-seq data analysis.

## Materials and methods

### Data Preprocessing

Any type of high-dimensional single-cell data can be de-sparsified using WIMC. However, before analysing the single-cell gene expression data, the raw scRNA-seq data often necessitates particular preprocessing and normalization to ensure that technical effect does not distort downstream analysis results. Given a raw data generated by sequencing machine, the quality control preformed based on following three criteria: 1) the number of counts per barcode (count depth), 2) the number of genes per barcode, and 3) the proportion of counts from mitochondrial genes per barcode. Since low-quality cells or empty droplets frequently have very few genes and a large number of detected genes may represent doublets, we begin by filtering cells with unique genes detected in excess of 3,000 or fewer than 200. Next, we filter out genes that are not expressed in fewer than 20 cells. In addition, cells with a mitochondrial count greater than 5 percent are removed, as cells with a relatively high mitochondrial count may be implicated in respiratory processes. We next perform a log normalization on the quality-controlled data matrix to reduce the variability of data before applying WIMC.

### Wavelet-based Imputation

The “dropout” events have an effect on the signal for each gene, which results in the loss of gene-gene relationships in the data. To overcome this sparsity, we develop wavelet-based imputation method, a non-parametric approach for recovering missing gene expression in scRNA-seq data. In our imputation method to de-sparsify sparse scRNA-seq data matrix (Fig.5A), the M-band DWT decomposes the count matrix denoted by s into M different frequency components, as shown in (3). In this paper, we select *M* = 2, 3, and 4, respectively, where the case of *M* = 2 corresponds to Daubechies wavelets family. The decomposition can be performed using the DWT matrix (Fig.5B). The DWT projects the data into orthogonal subspaces with different frequencies, allowing us to extract the information partially hidden by the canonical mixture data matrix.

**Fig 5.**
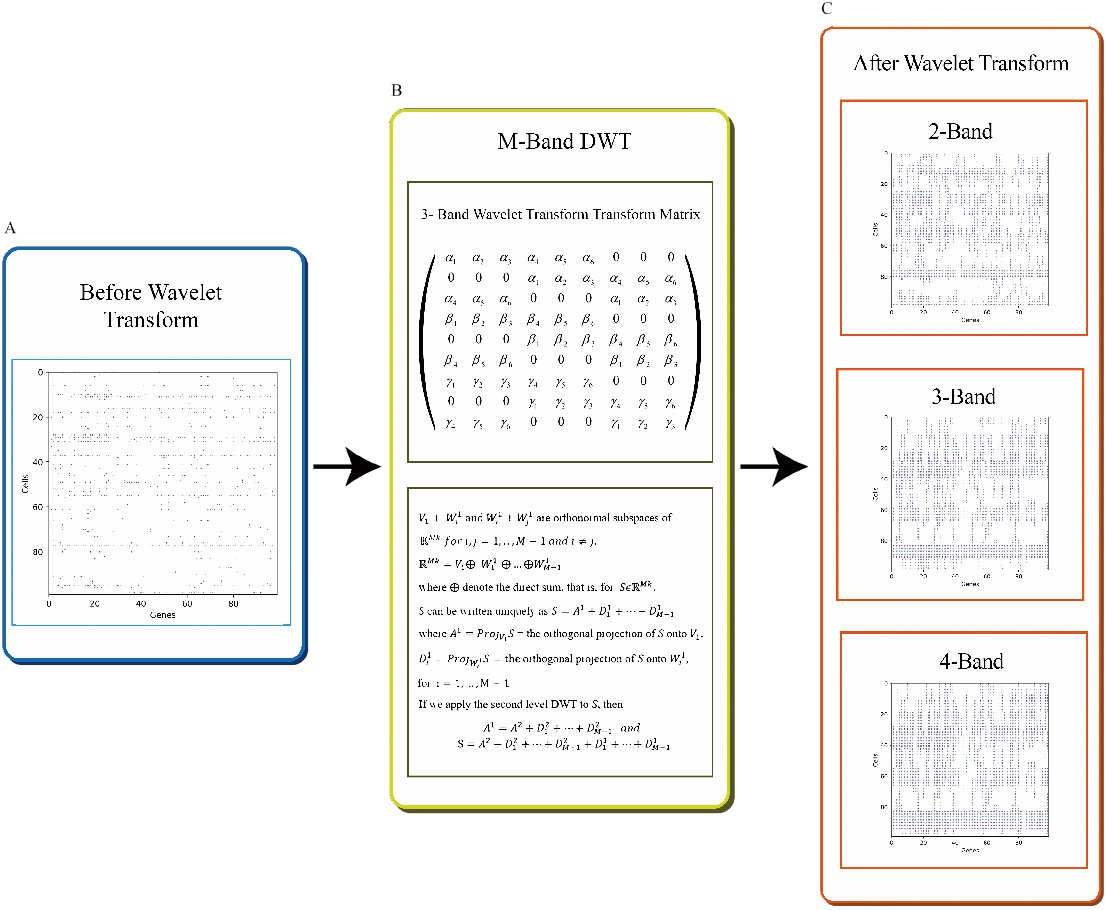
Demonstrating M-band orthogonal DWT imputation algorithm. (**A**) scRNA-seq sparse data matrix. (**B**) With applying M-band DWT to impute sparse matrix, the top panel shows a 3-band wavelet transform matrix while the bottom panel illustrates mathematical overview of M-band orthogonal DWT. (**C**) Resulting in imputed data matrices based on 2-band, 3-band and 4-band 2-regular DWT, successively.

The DWT can alternatively be viewed as a weighted moving average, with the weights functioning as corresponding filters. Consider the M-band R-regular low-frequency filter [*a*_1_, …, *a_k_*] transform the original data *S* into wavelet domain 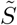 as following:

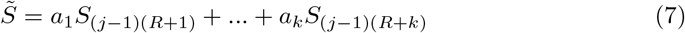

This process can de-sparsify the canoncial data matrix effectively, as the imputed data will usually be non-zero whenever at least one of its neighbors is non-zero in canonical data (Fig.5C).

Unlike traditional moving average method which uses the filter [1/*k*,…, 1/*k*] in (7), wavelet-based filters satisfy a set of orthogonal condition. Thus, the information in imputed data matrix under each component does not overlap theoretically. In addition, the DWT preserves the energy of the data since the *l*_2_-norm remains unchanged under orthonormal transform.

### Assessment using intersection analysis

We assess the performance of WIMC via computing the intersection of
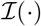, where 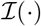 means the data conducted by PCA and UMAP. To examine the number of total clusters in multi-view windows, we conduct a test based on hyper-geometric distribution among clusters in different components. To be concise, : we describe the case of two components as follows. Suppose the low-frequency components are clustered as *L*_1_, …, *L_r_*, and *H*_1_,…, *H_s_* for high-frequency components. Denote *n_jk_* be the observed number of cells belonging to both *L_j_* and *H_k_*, and *n*_*j*+_ (*n*_+*k*_) be the total number of cells in *L_j_* (or in *H_k_*), respectively. we get a r by s contingency table, and our goal is to test the number of non-zero *n*_*jk*_’s. That is, to test

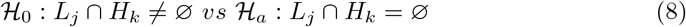

To consider the effect of possible random error in clustering algorithm, we first reject *H*_0_, for *n_jk_* < 3. Otherwise, if the cell clusters were distributed in chaos, then the expected number of *n_jk_* would be of hyper-geometric distribution as *n*_+*k*_ draws among a size-N population containing *n*_*j*+_ target objects. We calculate the p-value as the probability that less than *n_jk_* target objects are drawn, and set significant level 0.01.

Furthermore, we calculate the distribution of significant genes among different frequencies components. Specifically, let *G*_0_ be the set of significant genes in the raw data, *G*_1_ be that in the low-frequency component, and *G*_2_,…, *G_m_* be that in the high-frequency components. For each non-empty subset, Λ ∈ {0, 1,…, *m*}, we calculate 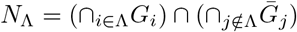 i.e., the set of genes in the selected components but not in the other components. If |Λ| = 1, |*N*λ| denotes the number of exclusive genes in the corresponding component. Note that the components of different frequencies under the DWT are orthogonal to each other, if |Λ| ≥ 2 and 0 ∉ Λ, only a few genes will contain in such *N*_λ_ theoretically. The smaller number of |*N*_λ_| for such intersection yields a better resolution of clustering.

## Supporting information

Detailed mathematical theory for DWT and results for more datasets are displayed in supplement materials.

## Data Availability

The breast cancer datasets, including CID-3921, CID-4495, CID-4523 and CID-4463 are available from the Gene Expression Omnibus (GEO) database with accession numbers GSM5354515, GSM5354530, GSM5354536 and GSM5354527, respectively. The Colon cancer dataset is available from GEO with accession number GSM4143678, while Peripheral Blood Mononuclear Cells dataset is accessible from https://satijalab.org/seurat/articles/pbmc3k_tutorial.html.

## Code Availability

The code for WIMC can be found at https://github.com/wangsresearchgroup/WIMC.

